# Dynamics of Plasma Lipidome in Progression to Islet Autoimmunity and Type 1 Diabetes – Type 1 Diabetes Prediction and Prevention Study (DIPP)

**DOI:** 10.1101/294033

**Authors:** Santosh Lamichhane, Linda Ahonen, Thomas Sparholt Dyrlund, Esko Kemppainen, Heli Siljander, Heikki Hyöty, Jorma Ilonen, Jorma Toppari, Riitta Veijola, Tuulia Hyötyläinen, Mikael Knip, Matej Oresic

**Author notes:** Corresponding and shared senior authors: Mikael Knip, M.D., Ph.D.; Children’s Hospital, University of Helsinki, P.O.Box 22, FI-00014 Helsinki, Finland.; Matej Orešič, Ph.D. Turku Centre for Biotechnology, University of Turku and Åbo Akademi University, Tykistokatu 6, FI-20520 Turku, Finland.

## Abstract

Type 1 diabetes (T1D) is one of the most prevalent autoimmune diseases among children in Western countries. Earlier metabolomics studies suggest that T1D is preceded by dysregulation of lipid metabolism. Here we used a lipidomics approach to analyze molecular lipids in a prospective series of 428 plasma samples from 40 children who progressed to T1D (PT1D), 40 children who developed at least a single islet autoantibody but did not progress to T1D during the follow-up (P1Ab) and 40 matched controls (CTR). Sphingomyelins were found to be persistently downregulated in PT1D when compared to the P1Ab and CTR groups. Triacylglycerols and phosphatidylcholines were mainly downregulated in PT1D as compared to P1Ab at the age of 3 months. Our study suggests that children who progressed to islet autoimmunity or overt T1D are characterized by distinct lipidomic signatures, which may be helpful in the identification of at-risk children before the initiation of autoimmunity.

## Introduction

Type 1 diabetes mellitus (T1D) is a heterogeneous, chronic autoimmune disorder characterized by selective loss of the insulin-producing pancreatic beta cells^1^. Over the past decades, the incidence and prevalence of T1D has notably increased among the children in Western countries^2^ and the number of cases is projected to double between 2005 and 2020 among European children^3^. However, no preventive strategies exist and the underlying pathogenesis of T1D is poorly understood.

Human leukocyte antigen (HLA)-conferred genetic susceptibility remains by far the strongest genetic risk factor for the disease^4^, yet less than 10% of the individuals with the high-risk HLA class II genotype progress to T1D^5^. This suggests that extrinsic factors probably contribute to disease development and progression either by triggering the immune response or by modifying the destructive process of beta cells in genetically predisposed individuals. Substantial evidence suggests that the first signs of beta cell autoimmunity may be initiated during the first year of life^6^ and a large body of literature^1, 2, 5, 7^ indicates that environmental factors and gut microbes play an important role in the disease development.

Determination of islet autoantibodies is the best known approach for identifying individuals at markedly increased risk for developing T1D before the onset of clinical symptoms^8^. Approximately 15% of children who progress to a single autoantibody develop T1D later in life, while for those who progress to multiple autoantibodies the risk is substantially higher (84% after-follow-up for 15 years)^9^. Earlier metabolomics studies also imply that children who progress to T1D are characterized by a specific metabolic profile which can be detected already prior to the appearance of islet autoantibodies^10, 11, 12^, suggesting that there is a link between the immune system, host metabolism and the development of T1D. However, the information whether the observed lipid-related changes are specifically associated with progression to T1D or with islet autoimmunity *per se* is still scarce. Here we analyzed plasma lipidome in a prospective series of samples which included children who progressed to overt T1D and control children who did not develop T1D or islet autoantibodies. We also included children who progressed to sero-positivity for at least a single islet autoantibody but did not progress to T1D during the follow-up, i.e., the at-risk children who will most likely not develop T1D^9^.

## Results

### Age related changes in the lipidome

We analyzed plasma lipidomics prospectively in three study groups: 40 progressors to T1D (PT1D), 40 who progressed to at least a single Ab but not to T1D during the follow-up (P1Ab), and 40 controls (CTRL) subjects who remained islet autoantibody negative during the follow-up until the age of 15. For each child, up to six time points were included in the analysis, corresponding to the ages of 3, 6, 12, 18, 24 or 36 months (**Fig. 1**). None of the children were diagnosed with T1D at the time of sample withdrawal. The lipidomics dataset (n =428) used for the analysis included identified lipids from the following lipid classes: cholesterol esters (CE), diacylglycerols (DG), lysophosphatidylcholines (LPC), phosphatidylcholines (PC), phosphatidylethanolamines (PE), sphingomyelins (SM) and triacylglycerols (TG).

**Figure 1.**
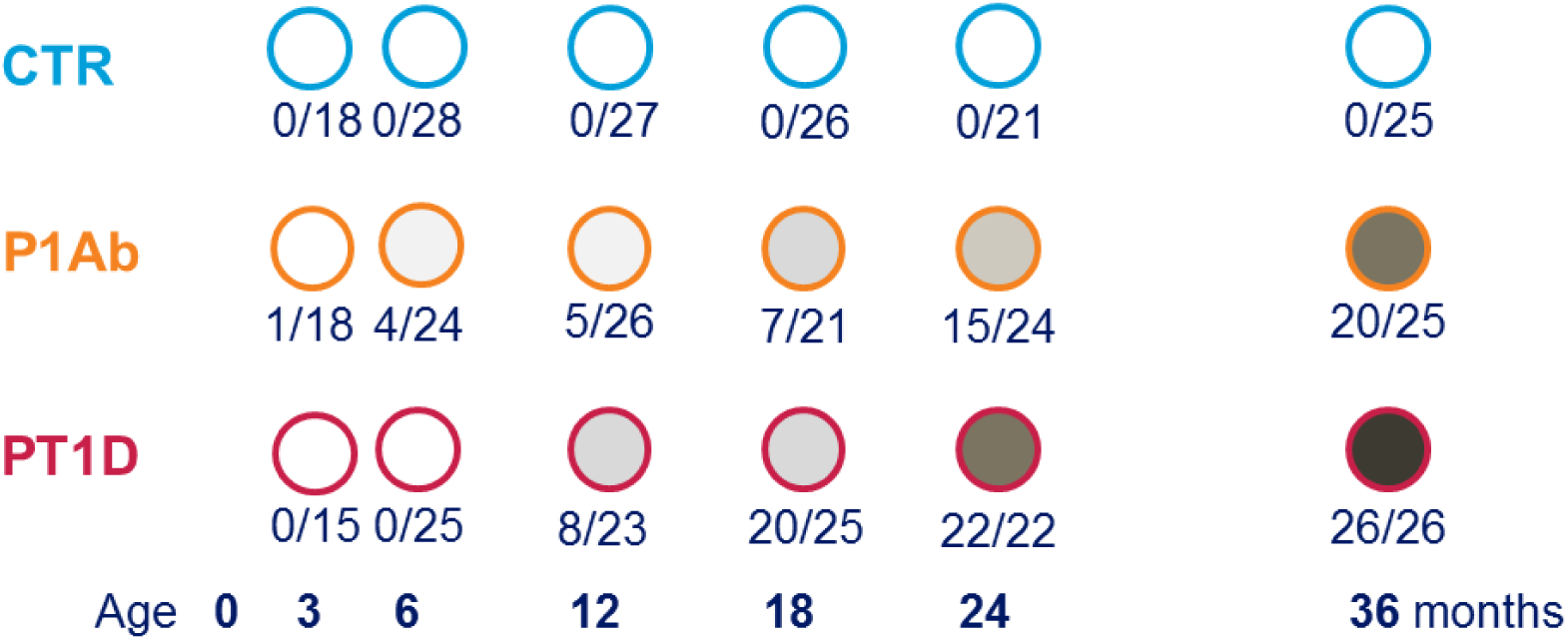
An overview of sample and the study design. The study cohort comprises the samples from children, who progressed to T1D (PT1D), who developed at least a single antibody (Ab) but not T1D during the follow-up (P1Ab), and control (CTRL) subjects who remained islet autoantibody negative during the follow-up until the age of 15 years. For each child, longitudinal samples were obtained corresponding to either of the ages of 3, 6, 12, 18, 24, and 36 months. In this figure the numbers of Ab positive samples at any given time point (X) and the numbers of samples at that time point (Y) is depicted in format (X/Y) for the specific age cohort.

Principal Components Analysis (PCA)^13^ of the preprocessed lipidomics data revealed a clear age-related trend. In order to specifically determine the age-related associations in the plasma lipidome, analysis of variance (ANOVA)-simultaneous component analysis (ASCA)^14^ was performed with factors age, gender and case status (PT1D, P1Ab, CTR) including their interactions separating the variation into contributions from the different factors. Age had the strongest effect (6.01, p = 0.0010) on the plasma lipidomic profile when compared to the case group (0.89, p = 0.0050) and gender (0.60, p = 0.0020). However, the interactions between these factors remained insignificant. The score plot in **Fig. 2a** highlights the age-related trend and the corresponding loadings plot (**Fig. 2b**) explains the class-specific lipid alteration related to age.

**Figure 2.**
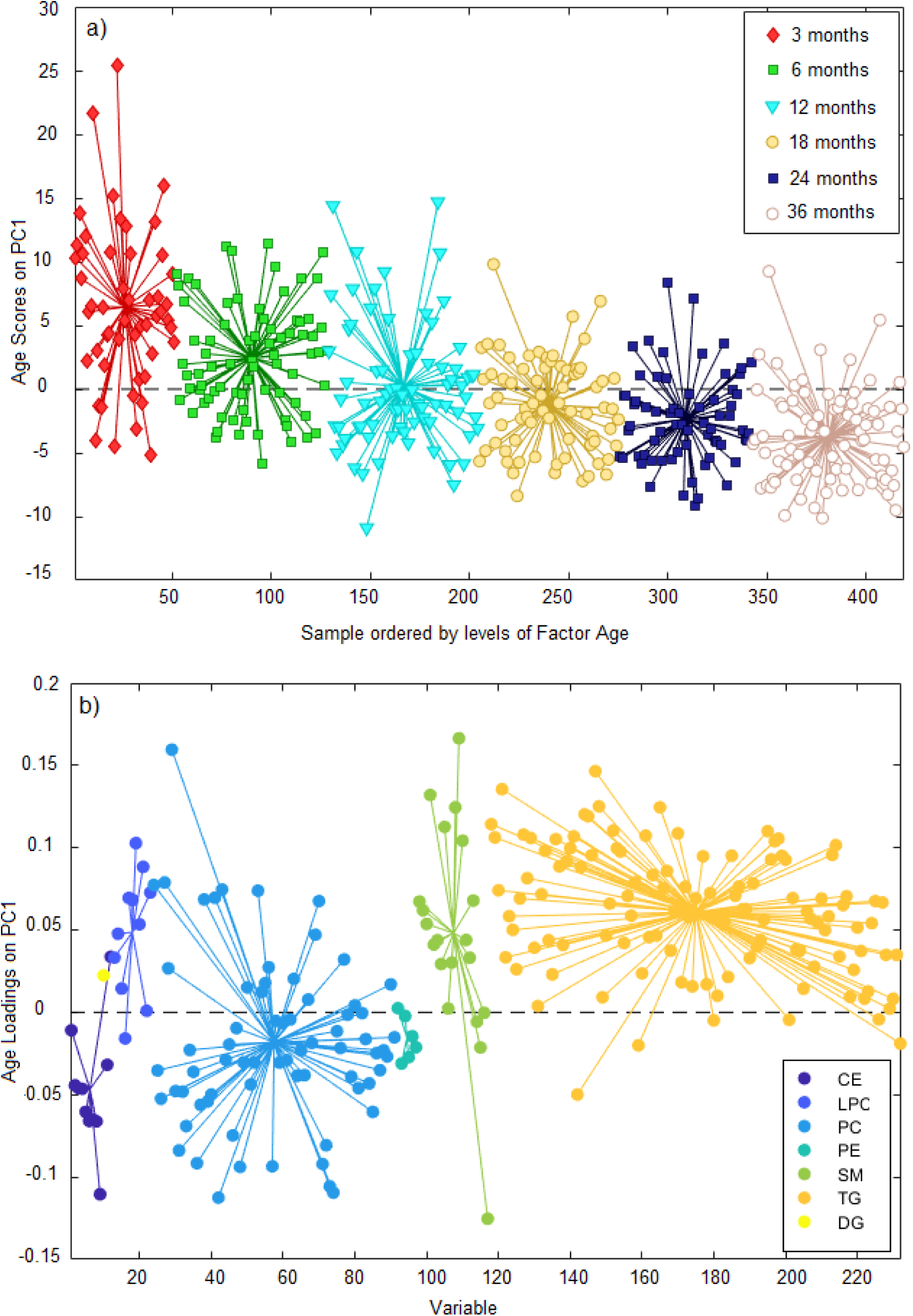
PCA score plots and loadings based on ASCA. (**a**) PC1 score plot based on ASCA. These scores represent the lipidomics dataset arranged according to age in the PCA score plot. Here each sample is represented by a point and colored according to the age. The age of the participants are marked on the x-axis while y-axis represents the sample score. Samples with similar score are clustered together. (**b**) The corresponding PC1 loading plot. The loadings explain the pattern seen in the score plot which provides the means to interpret the class specific lipid alteration related to age. X-axis is the variable order (i.e. lipids) and y-axis represents the lipids pattern corresponding to the score plot. Here, the levels of SMs and LPCs were elevated at early time points (3, 6, 12 months) while CEs and PEs appeared to be higher in later time point (18, 24, 36 months). Abbreviations: cholesterol ester (CE), diacylglycerol (DG), lysophosphatidylcholine (LPC), phosphatidylcholine (PC), phosphatidylethanolamine (PE), sphingomyelin (SM) and triacylglycerol (TG).

Specifically, the loadings depicts that the levels of SMs and LPCs were elevated at the early time points (3, 6, 12 months) while CEs and PEs appeared to be higher at later time points (18, 24, 36 months). In contrast, the levels of TGs and PCs varied over time with no clear age-related patterns.

### Serum lipidome in development of autoimmunity and progression to T1D

In order to minimize the confounding effect of age, data were analyzed separately in different age cohorts. Partial least squares discriminant analysis (PLS-DA)^15^ was performed to examine the differences between the three sample groups using combinations of various categorical variables (e.g. PT1D vs. P1Ab, PT1D vs. CTR) in each age cohort.

**Fig. 3** shows the VIP (variable importance for projection) dependent regression coefficient plot obtained from the PLS-DA analysis between PT1D (n=15) and CTR (n=18) at 3 months of age (AUROC = 0.78). The result show lower levels of SMs and higher levels of multiple PCs in children who later progressed to T1D as compared to CTR. Among the PCs, particularly those with a higher number of double bonds (>1) showed an increase in PT1D as compared to CTR (**Supplementary Fig. 1**). In addition, CE (20:4) and PE (36:4) appeared to be higher in PT1D, but no clear patterns were seen for the TGs and LPCs. For example, LPC (16:1) was elevated while LPC (16:0e) was downregulated in PT1D. Based on the VIP scores, SM (d36:2), SM (d36:1), SM (d34:1), SM (d38:2) and PC (40:4), PC (O-34:2), PC (34:2) and PC (O-36:2) were top ranked discriminating lipids in the supervised analysis. Univariate analysis showed that at the age of 3 months, a total of 12 lipids were altered between PT1D and CTR according to the nominal p-value < 0.05. These lipids included four PCs, seven SMs and one TG (**Supplementary Table 1**). However, none of these lipids passed significance at the selected false discovery rate (FDR) threshold of 0.1. The adjusted p-values for SM (d36:1), SM (d34:1), and SM (d36:2) were 0.15, 0.15, and 0.20, respectively.

**Figure 3.**
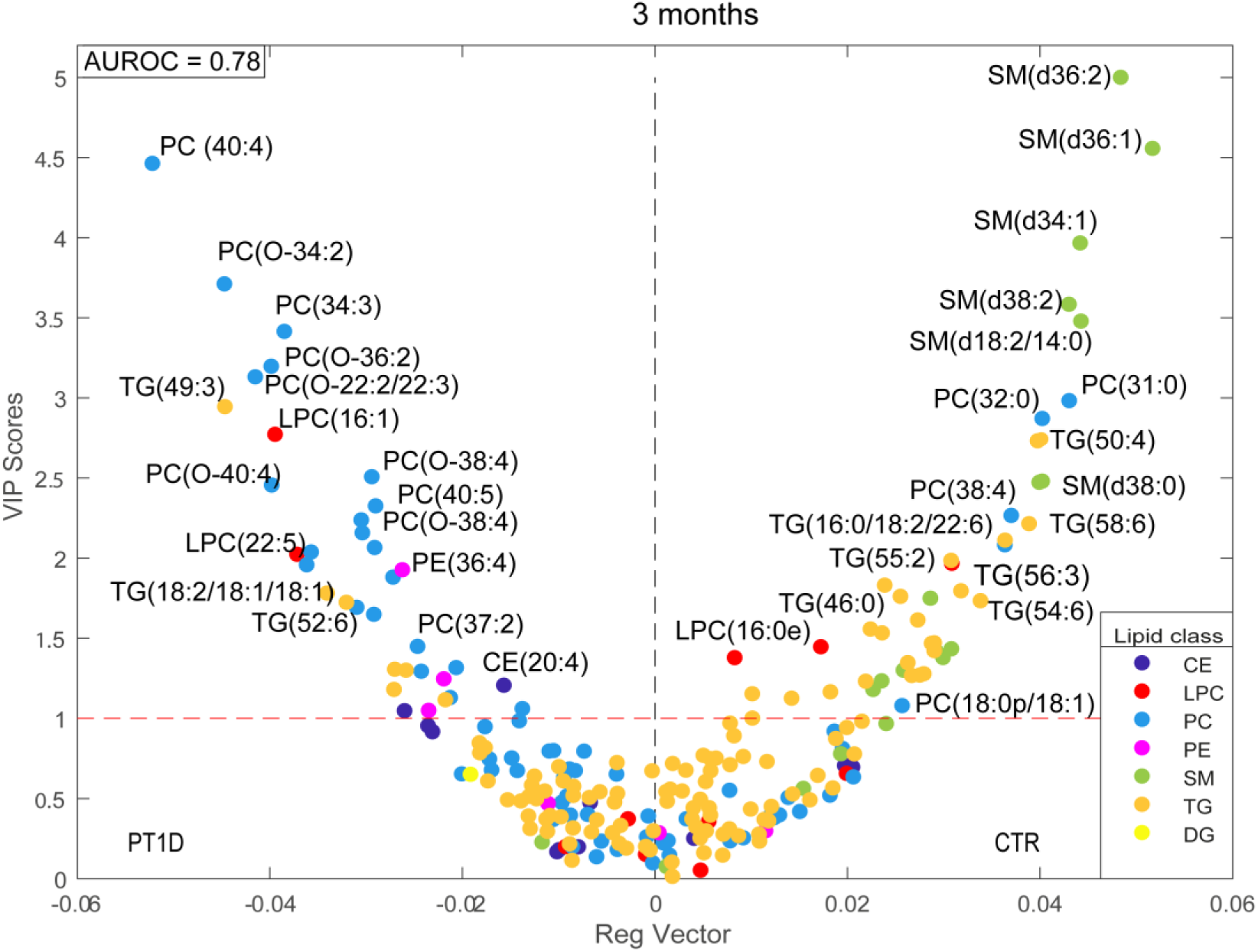
Regression coefficient plot from PLS-DA analysis. This plot has the regression coefficient on the x-axis and VIP scores in the y-axis. The positive and negative regression coefficients represent CTR and PT1D groups, respectively. Each lipid class is color coded. Abbreviations: cholesterol ester (CE), diacylglycerol (DG), lysophosphatidylcholine (LPC), phosphatidylcholine (PC), phosphatidylethanolamine (PE), sphingomyelin (SM) and triacylglycerol (TG).

PLS-DA analysis comparing PT1D (n=15) and P1Ab (n=18) at the age of 3 months revealed higher levels of SMs and TGs in the P1Ab group (AUROC = 0.67; **Supplementary Fig. 2a**). Univariate analysis showed that a total of 45 lipids being different between PT1D and P1Ab (nominal p-value<0.05). These lipids included one LPC, 12 PCs, one PE, nine SMs and 22 TGs. All of these lipids passed the FDR threshold of 0.1 (**Fig. 4a**, **Supplementary Table** 2). With the exception of LPC (16:0), PC (O-22:2/22:3), PC (O-36:2), PC (O-40:5), most of the lipids were decreased in PT1D group (**Fig. 4a**). However, no clear pattern with respect to the acyl chain composition was observed in any specific lipid class.

**Figure 4.**
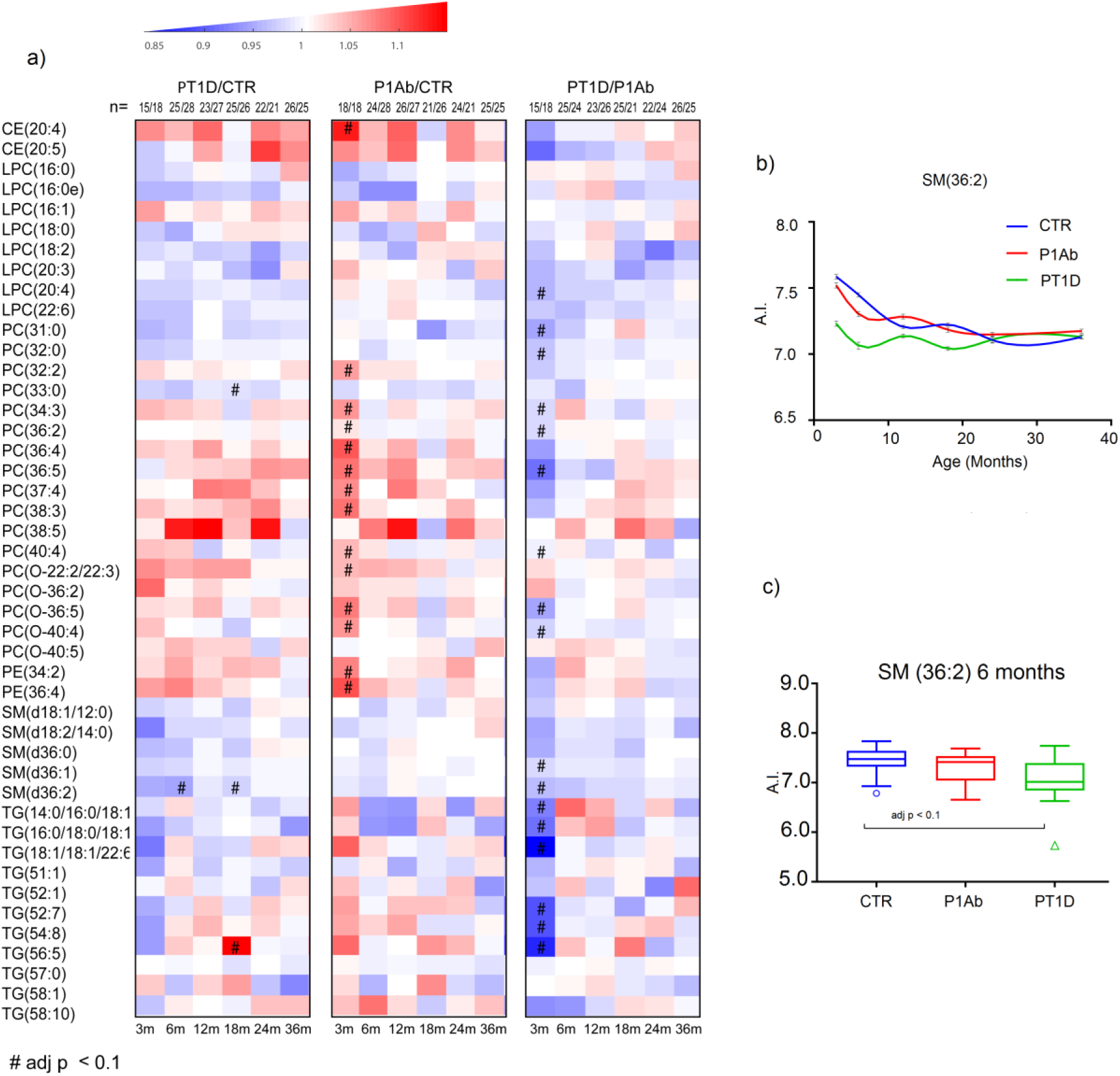
The differences in global serum lipidome between the three study groups. (**a**) Heat map showing 45 lipid species representative of lipid classes that change between PT1D, P1Ab and CTR. Differences in lipid concentrations were calculated by dividing mean concentration in PT1D by the mean concentrations in P1Ab and CTR. Differences were determined at 3, 6, 12, 18, 24 and 36 months of age between the study groups. (**b**) The smoothened line plot based on the mean SM (36:2) concentration in the three study groups. The error bar shows the standard error of the mean. (**c**) Concentrations of SM (36:2) in the three study groups at 6 months of age. # represents the adjusted p-values < 0.1.

Next, PLS-DA analysis between P1Ab (n=18) and CTR (n=18) at 3 months of age revealed increased levels of PCs and PEs in the P1Ab group (AUROC = 0.65; **Supplementary Fig. 2b**). However, no clear patterns were observed for the other lipid classes. Analysis of individual lipids revealed that a total of 45 lipids were altered between the two study groups (nominal p-value <0.05). These lipids included two CEs, 38 PCs, four PEs, and two TGs. Fourty-four of these lipids passed the selected FDR threshold of 0.1 (**Supplementary Table 3**). The heatmap highlights that most of the PCs, with the exception of PC (33:0) were increased in the P1Ab group. ASCA analysis also revealed that the study group had a significant effect (p = 0.022) on the plasma lipidome profile of the children while the effect of gender (p = 0.60) and its interaction (p = 0.95) remained insignificant in the 3-month cohort.

Similarly, we analyzed the lipid concentration differences between the three study groups in the remaining longitudinal series at 6, 12, 18, 24 and 36 months. There was no persistent trend with respect of lipid differences between the groups (**Fig. 4a**), with the exception for SMs which were persistently low in PT1D as compared to P1Ab and CTR (**Fig. 4b**). Low levels of SMs were particularly observed at 6, 12 and 18 months (**Fig. 4b**). However, only one of the SMs, SM (d36:2), differed significantly (adjusted p < 0.1) between the PT1D and CTR at 6 and 18 months of age (**Fig. 4c)**. In addition, PC (33:0), TG (56:5) and TG (18:2/18:1/18:1) passed the FDR threshold of 0.1 at 18 months of age (**Fig. 4a**).

### Lipidome in progression to type 1 diabetes

Next, we sought to determine the total lipid class specific changes in the longitudinal series of samples. Individual lipid concentrations in each lipid class were added, then subsequent data analysis considering each lipid class as a variable was performed. We found clear alterations in phospholipids, in particular SMs were significantly reduced in the PT1D group at the ages of 3 and 18 months when compared to the P1Ab group (**Fig. 5a**) and at the age of 3 month when compared to CTR (**Fig. 5b**). TGs and PCs were significantly lower in PT1D than in P1Ab at the age of 3 months (**Fig. 5a & 5c**). PEs and PCs were also higher in P1Ab than in CTR at the age of 3 months.

**Figure 5.**
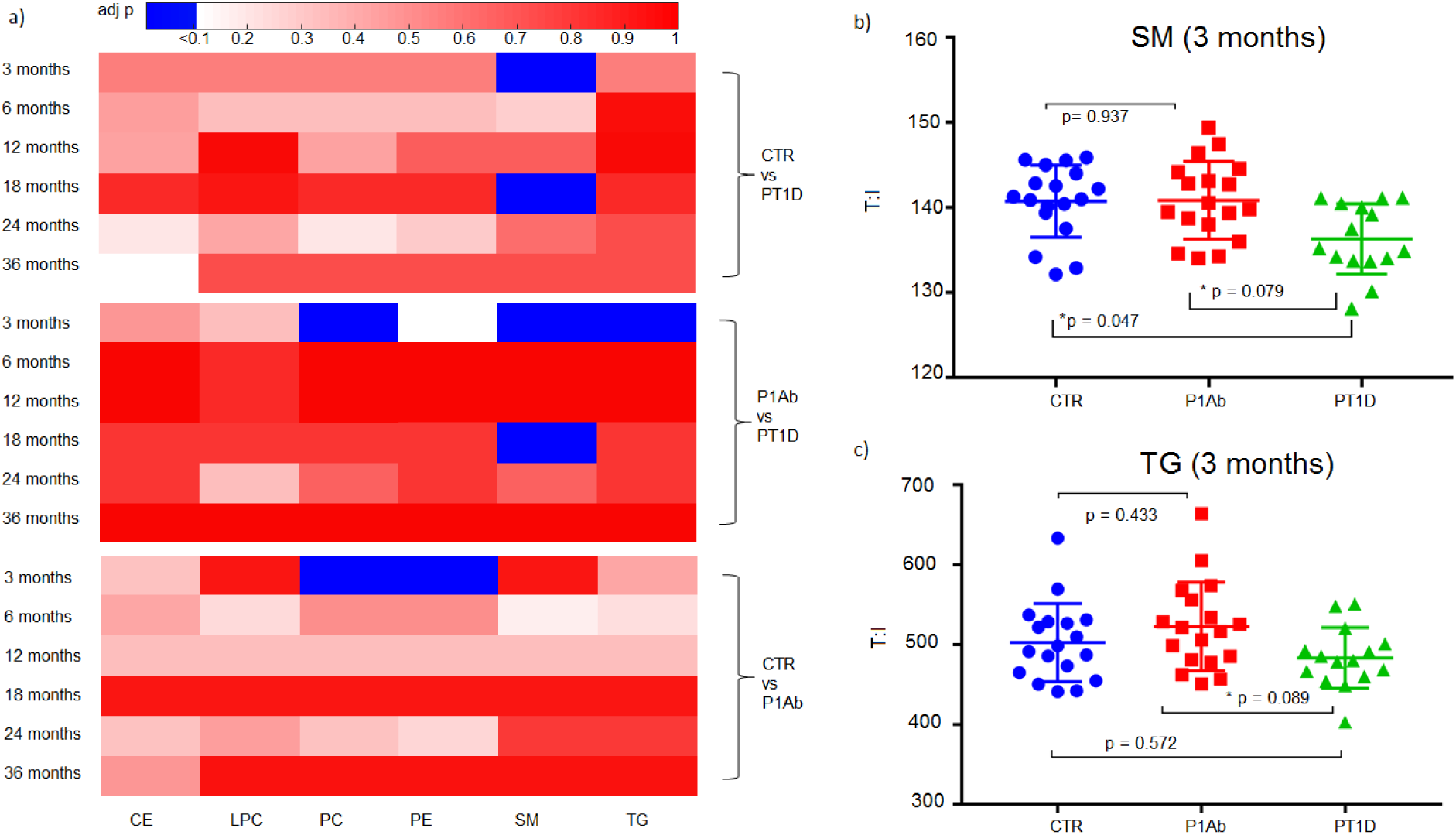
Total lipid concentration in each lipid class in the longitudinal cohort at the ages of 3, 6, 12, 18, 24 and 36 months. Adjusted p-values < 0.1 are colored blue. SMs were significantly downregulated in the PT1D group at the age of 3 months compared to P1Ab and CTR (adjusted p = 0.047). TGs were also significantly downregulated in PT1D as compared to P1Ab (adjusted p = 0.079) at the age of 3 months.

### Lipidome changes before the emergence of islet autoantibodies

In order to examine whether the circulating lipids were associated with the appearance of islet autoantibodies, the plasma lipidome in PT1D with one Ab (n= 21) and P1Ab (n=27) groups were compared between the samples obtained before and after the appearance of the first islet autoantibody, respectively. We applied the multi-level approach to exploit the paired data structure. Multi-level analysis revealed several differences before and after the appearance of islet autoantibodies. Specifically, SMs (SM (d36:1), SM (d36:2)) and LPCs (LPC (18:0), LPC (18:2), LPC (16:0)) were higher while CEs (CE (20:4), CE (20:5)) were lower before seroconversion than after it in both PT1D and P1Ab (**Fig. 6**). As an exception to this pattern, LPC (20:3) appeared to be elevated after the emergence of the first autoantibody in P1Ab. Next, we set out to examine the individual lipid profiles by univariate analysis. The pairwise analysis revealed a total of 27 (two CEs, three LPCs, 11PCs, five SMs, six TGs) and 16 (two CEs, two LPCs, six PCs, four SMs, two TGs) lipids, respectively, being altered in both study groups (nominal p-value <0.05, **Supplementary Table 4-5**). However, only CE (20:5) passed the level of significance at the selected FDR threshold of 0.1 in P1Ab.

**Figure 6.**
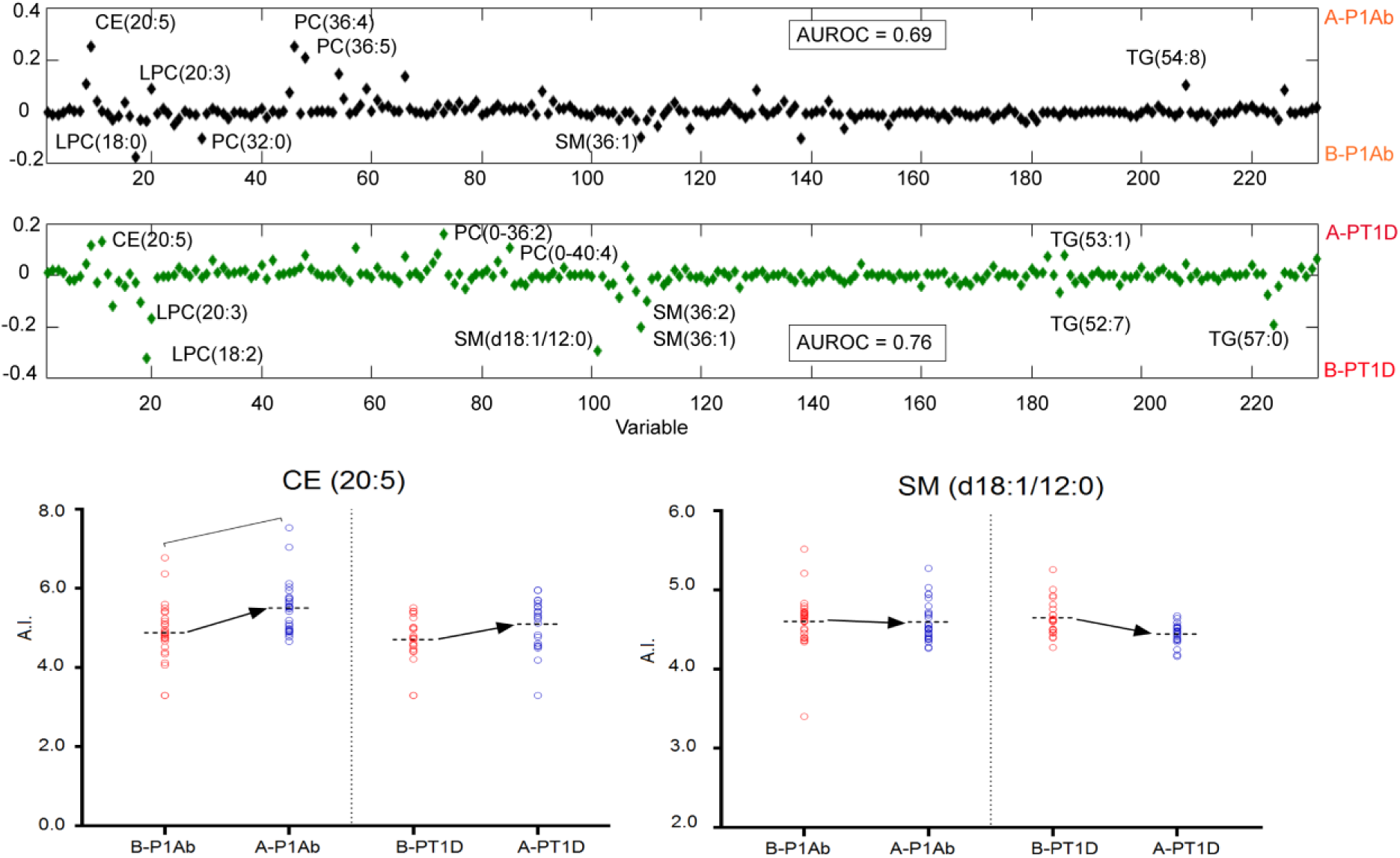
The ML-PLSDA regression coefficient plot showing the most discriminating lipids between the samples obtained before the first islet autoantibody appeared and the samples after the emergence of the first islet autoantibody in PT1D and P1Ab groups. The plot shows that SM (d36:1), SM (d36:2), LPC (18:0), LPC (18:2) and LPC (16:0) were up-regulated while CEs (CE 20:4), CE (20:5)) were downregulated before the seroconversion. The pairwise scatter plot shows that SM (18:1/12:0) and CE (20:5) are differentially altered in P1Ab as compared to PT1D. However, only CE (20:5) passed the FDR threshold of 0.1 in P1Ab after seroconversion. Abbreviations: Before seroconversion in P1Ab (B-P1Ab), after seroconversion in P1Ab (A-P1Ab), before seroconversion in progressors (B-PT1D), after seroconversion in progressors (A-PT1D).

We also studied whether lipidome profiles associated with the appearance of specific islet autoantibodies. The plasma lipidome were compared before and after the emergence of first insulin autoantibodies (IAA) in P1Ab and PT1D with one Ab. Pairwise comparisons were also performed by combining other autoantibodies as a group (islet cell antibodies (ICA), islet antigen 2 autoantibodies (IA-2A), and GAD autoantibodies (GADA) in the remaining children. The pairwise comparison revealed a total of 17 (in PT1D with first IAA), one (in PT1D with all others autoantibodies excluding IAA), 22 (in P1Ab with first IAA), and 11 (in P1Ab with all others autoantibodies excluding IAA) lipids being altered at a nominal p-value <0.05 before and after the appearance of specific islet seroconversion (**Supplementary Tables 6-9**). However, none of the lipid species passed the significance at the FDR threshold of 0.1.

Next, we analyzed the lipid concentration differences between the specific islet (IAA vs others including IA-2, GADA, ICA) autoantibodies study groups in the longitudinal series at 3, 6, 12, 18, 24 and 36 months. There were no significant (FDR threshold of 0.1) lipid differences between the specific islet groups, with the exception for 3 months of age, where we found a total of 23 lipids (two CEs, one PC, seven SMs, 13 TGs) being altered when comparing PT1D with IAA to P1Ab with IAA (**Supplementary Table 10**). All of these lipids passed significance at the FDR threshold of 0.1. SMs, TGs and PC were lower while CEs appeared to be higher at the age of 3 months in the PT1D group with IAA than in the P1Ab group with IAA as the first autoantibody (**Supplementary Fig. 3**).

## Discussion

Here we showed that specific lipid changes precede islet autoimmunity and T1D. Our results revealed that the children who progress to T1D in the follow-up tend to have a distinct and persistently dyregulated lipid profile as compared to those who later progress to islet autoimmunity but not to T1D. Our observations also show that plasma lipids, in particular CEs, PEs, SMs and LPCs, are strongly affected by age during the early development. This is in agreement with earlier studies which suggest that gender, age, and body mass index have significant impact on the longitudinal trajectories of plasma lipids^16, 17^. Several studies depict that serum total cholesterol and TGs gradually increase with age during the early development^18, 19^, but the data specifically from infants are limited to phospholipids such as SMs and LPCs^20^. Our findings are not unexpected however, because various host related and environmental factors including dietary changes during the development can impact the lipid profiles over time^21^.

We found that differences in the lipidomes between the three study groups were most pronounced at the earliest age of 3 months. SM levels were low in the PT1D group at the ages of 3 and 18 months. At the level of individual lipid species, only SM (d36:2) differed between PT1D and CTR at 6 and 18 months of age, whereas this lipid also differed between PT1D and P1Ab at 3 months of age. Although not significant statistically, most of the other SMs species tend to be persistently lower in PT1D than in both CTR and P1Ab. Interestingly, the lipid-related associations with progression to T1D appear to be the strongest in the group of children who first developed positivity for IAA, which are also the most common first autoantibody detected in young children who later progress to T1D^5^ and tends to be associated with higher risk of progression as compared to other autoantibodies^22, 23^. Previous studies by us and others have also reported decreased serum SM levels in children who later progressed to T1D as well as in children with newly diagnosed T1D^24, 25^. Sphingolipids are among the most abundant choline containing phospholipids in the mammalian cells, tissues and in the circulation. Sphingolipids have been shown to be potent regulators of immune cell activity, particularly in the context of T-cell-mediated immunity^26^. In addition, sphingolipids are crucial for plasma membrane organization and thus their concentrations might affect multiple T-cell processes such as activation, expansion, cell-cell interactions and homing into secondary lymphoid tissues that play an important role in immunity, inflammation and inflammatory disorders^27^. Much of this regulation has been demonstrated by modulating the activity of the sphingomyelinase family of enzymes which catalyze the breakdown of sphingomyelin into ceramide and other precursor metabolites^26^. Emerging evidence suggests that dysregulated sphingomyelinase activity in conjunction with cellular apoptosis can result in chronic inflammatory disorder^28^. Furthermore, deficiency for the acid sphingomyelinase has been demonstrated to enhance the proliferation of immunosuppressive T regulatory cells and inhibition of brain cell death in mice^29^. Intriguingly, disruption of sphingolipid metabolism has also been increasingly associated with the development and progression of other metabolic conditions, including obesity and type 2 diabetes^30^. Our results suggest that SM metabolism is disrupted among the T1D progressors from early age, already preceding the appearance of islet autoantibodies.

While confirming the earlier findings that progression to T1D is associated with decreased concentrations of SMs^24^, the current study also shows that these SM-related changes are not associated with progression to islet autoimmunity *per se*, thereby, suggesting that deregulated sphingolipid metabolic pathway might be a possible contributor to the T1D etiology. Apart from the sphingolipid metabolic pathway, defect in T-cell derived exosomes cargo could be linked with the downregulated sphingolipids observed in P1TD^31^. Exosomes are small (40–150 nm) vesicles released from cells that carry the host cell-derived cargo such as proteins, lipids, and RNA. T-cell derived exosomes are highly enriched in sphingomyelin content compared to the human T lymphocyte cell line (Jurkat T cells)^32^. Insulinoma-derived exosome have been reported to activate T cells in the non-obese pre-diabetic mice and to accelerate insulinitis in the non-obese diabetes-resistant mice^31^, implicating a role for aberrant exosome signaling in the pathogenesis of autoimmune diabetes.

An important consideration in the search of metabolic biomarkers of T1D progression is that blood in young infants is sampled in non-fasting state. In the earlier study, SMs were found to be least variable within-individuals in nonfasted serum samples^12^. This suggests that SMs may also hold promise as predictive biomarkers of T1D and thus complementing genetic testing and islet autoantibody determination. Clearly more and larger studies in heterogeneous populations are needed to examine the complementary diagnostic value of SMs.

The total TGs and PCs were downregulated in PT1D as compared to P1Ab at 3 months of age. Consistent with these observations, other studies have reported perturbations in phospholipids during the early infancy among the T1D progressors^10, 24, 25^ as well as decreased TGs in newborn infants who later progressed to T1D^24^. Choline regulates secretion of triglyceride rich lipoprotein particles and its deficiency has been linked to lower serum TG concentration^33^. Since PCs are a major source of choline; an important epigenetic regulator in the body^34^, it is conceivable that alterations in the choline metabolism during infancy, e.g. by diet, may affect the PC levels, resulting in the decreased levels of TGs as well as PCs. In addition, the decreased levels of TGs may also suggest increased energy demand in PT1D as compared to P1Ab^35^. TGs are concentrated stores of metabolic energy in biological tissues^36^. During energy demand, TGs are hydrolyzed into free fatty acids that are taken up by other tissues or cells to meet the energy requirements of the organism^37^. Intriguingly, the energy demand of regulatory T cells is extraordinary and strongly favors fatty acid oxidation^38^. Compelling evidence shows that regulatory T cells are altered in T1D^39, 40^ and they play an important role in the stabilization of immune responses^41^. Downregulated TGs as detected in our studies thus suggest distinct energy linked metabolic state in PT1D as compared to P1Ab that might have a direct effect on regulatory T-cell function.

The appearance of autoantibodies was associated with down-regulation of SMs and LPCs and up-regulation of CEs. LPC is the main phospholipid component of oxidized low density lipoprotein (oxLDL) produced as a hydrolysis product of PC^42^. LPC regulates various intracellular signaling events triggered by the oxLDL, including modulation of the toll like receptor (TLR) signaling pathway and activation of macrophages which are the hallmarks of the inflammatory and immunological processes^42, 43^. A growing body of evidence suggests that LPCs have a functional role in the etiology of inflammatory diseases such as atherosclerosis ^44^ and multiple sclerosis^45^. Our findings are in line with earlier observations that LPC dysregulation precedes the appearance of islet autoantibodies^24^.

Up-regulation of CEs following seroconversion to positivity for islet autoantibodies is a novel finding. Previous studies have provided evidence for the inflammatory effects of cholesterol accumulation^46^. Cholesterol accumulation activated the enzyme lecithin cholesterol acyltransferase (LCAT) that modifies cholesterol to CE. The esterified cholesterols are subsequently taken up by the liver and discharged through bile excretion^47^. An efficient cholesterol esterification would improve the reverse cholesterol transport leading to a reduction in inflammatory lesions^48^. In contrast, reduced cholesterol efflux may trigger pro-inflammatory signaling pathways such as TLR signaling, which results in the modulation of inflammatory responses^49^. Intriguingly, impaired cholesterol efflux has been associated with the pathogenesis of various inflammatory disease including atherosclerosis, rheumatoid arthritis and systemic lupus erythematosus^50, 51^. Given the abnormality observed in other autoimmune pathologies, it is possible that increased CE accumulation after seroconversion in islet autoimmunity, particularly in P1Ab can be interpreted as a sign of a protective mechanism. Meanwhile, less CEs efflux in the PT1D group might be an indication of failing immune protection. However, understanding of the role of cholesterol homeostasis with respect to islet autoimmunity is currently limited.

Taken together, our findings support earlier studies by us and others, which suggest that dysregulation of lipid metabolism precedes islet autoimmunity and T1D, as well as add a new knowledge that the observed changes are specifically and persistently associated with progression to overt disease. The distinct lipidomics profile which is associated with the progression to T1D appears to be most pronounced in very young children, i.e. at 3 months of age in our study. This suggests (1) that diversification of dietary patterns already during the development may mask some of the T1D-associated metabolic signatures and (2) that metabolite-based prediction of T1D may be most feasible in early life, already prior to the appearance of islet autoantibodies. Already in young infants, the lipidomics signatures of children who progress to T1D later in life are clearly different from those that do seroconvert to a single islet autoantibody but do not progress to overt disease. Since several lipids found dysregulated in the PT1D group displayed an opposite pattern in the P1Ab group in relation to CTR, our findings suggest that the observed metabotype may have not only a role in early T1D pathophysiology but may inversely also have a protective role in P1Ab. Since the current stratifications in studies aimed at T1D prevention are based on the detection of islet autoantibodies, lipidomics profiles may thus provide a valuable complementary tool for identifying children at highest risk of progression to T1D.

## Methods

### Sample and study design

The samples have been obtained from the Finnish Type 1 Diabetes Prevention and Prediction Study (DIPP) ^52^. The DIPP study has screened more than 220,000 newborn infants for HLA-conferred susceptibility to T1D in three university hospitals Turku, Tampere, and Oulu in Finland^53^. The subjects involved in the current study were chosen from the subset of DIPP children from the Tampere study center. Here, five longitudinal samples per child were collected between 1998 and 2012. For each child, longitudinal samples were obtained corresponding to either of the ages of 3, 6, 12, 18, 24, and 36 months. This cohort comprises the samples from 120 children: 40 progressors to T1D (PT1D), 40 who tested positive for at least one Ab in a minimum of two consecutive samples but did not progress to clinical T1D during the follow-up (P1Ab), and 40 controls (CTRL) subjects who remained islet autoantibody negative during the follow-up until the age of 15. The three study groups were matched by HLA-associated diabetes risk, gender and period of birth. Selected characteristics of the study subjects are shown in **Table 1**. In total 428 non-fasting blood samples were collected for this study. Plasma was separated within 30 minutes after the blood collection by centrifugation at 1600g for 20 minutes at room temperature. The plasma samples were stored at -80°C until analyzed.

**Table 1.** Demographic characteristics of study population.

|  | **PT1D** | **P1Ab** | **CTR** |
| --- | --- | --- | --- |
| **Gender (girls, boys)** | (14, 26) | (14, 26) | (14, 26) |
| **Age at time of diagnosis**<br>**(mean ± SD)** | 4.75 ± 2.94 | NA | NA |
| **Age at time of first seroconversion**<br>**(mean ± SD)** | 1.34 ± 0.58 | 3.05 ± 2.50 | NA |
| **HLA risk haplotypes** |  |  |  |
| **High risk (DR3 - DQA1\*05 - DQB1\*02)**<br>**/DRB1\*0401/2/4/5 - DQA1\*03 - DQB1\*0302, girls,**<br>**boys)** | (4, 5) | (1, 3) | (2, 4) |
| **Moderately increased risk\*** | (7, 14) | (9, 12) | (5, 9) |
| **Slightly increased risk\*** | (2, 8) | (1, 8) | (1, 6) |
| **Neutral\*** |  | (2, 1) | (3, 1) |
| **Decreased risk and/or not possible to form**<br>**haplotype\*** |  | (0, 2) | (3, 5) |

\*other than DR3 - DQA1\*05 - DQB1\*02) /DRB1\*0401/2/4/5 - DQA1\*03 - DQB1\*0302

### HLA genotyping

HLA-conferred susceptibility to T1D was analyzed using cord blood samples as described^54^. Briefly, the HLA-genotyping was performed with time-resolved fluorometry based assay for four alleles using lanthanide chelate labeled sequence specific oligonucleotide probes detecting DQB1*02, DQB1*03:01, DQB1*03:02, and DQB1*06:02/3 alleles^55^. The carriers of the genotype DQB1*02/DQB1*03:02 or DQB1*03:02/x genotypes (here x≠ DQB1*02, DQB1*03:01, DQB1*06:02, or DQB1*06:03 alleles) were categorized into the T1D risk group and recruited for the DIPP follow up program.

### Detection of islet autoantibodies

The participants with HLA-conferred genetic susceptibility were prospectively observed for the appearance of T1D associated autoantibodies (islet cell antibodies (ICA), insulin autoantibodies (IAA), islet antigen 2 autoantibodies (IA-2A), and GAD autoantibodies (GADA). These autoantibodies were measured in the Diabetes Research Laboratory, University of Oulu from the plasma samples taken at each follow-up visit as described ^56^. ICA were detected with the use of indirect immunofluorescence, whereas the other three autoantibodies were quantified with the use of specific radio binding assays^57^. We used cutoff limits for positivity of 2.5 Juvenile Diabetes Foundation (JDF) units for ICA, 3.48 relative units (RU) for IAA, 5.36 RU for GADA, and 0.43 RU for IA-2A. The disease sensitivity and specificity of the assay for ICA were 100% and 98%, respectively, in the fourth round of the international workshops on standardization of the ICA assay. According to the Diabetes Autoantibody Standardization Program (DASP) and Islet Autoantibody Standardization Program (IASP) workshop results in 2010–2015, disease sensitivities for the IAA, GADA and IA-2A radio binding assays were 36–62%, 64–88% and 62–72%, respectively. The corresponding disease specificities were 94–98%, 94–99% and 93– 100%, respectively.

### Analysis of molecular lipids

A total of 428 serum samples were randomized and extracted using a modified version of the previously published Folch procedure^58^. 10 µL of 0.9% NaCl, 40 µL of CHCl_3_:MeOH (2:1, v/v) and 80 µL of the 3.5 µg mL^-1^ internal standards solution (for quality control and normalisation purposes) were added to 10 µL of each plasma sample. The standard solution contained the following compounds: 1,2-diheptadecanoyl-*sn*-glycero-3-phosphoethanolamine (PE(17:0/17:0)), N-heptadecanoyl-D-*erythro*-sphingosylphosphorylcholine (SM(d18:1/17:0)),N-heptadecanoyl-D-*erythro*-sphingosine (Cer(d18:1/17:0)),1,2-diheptadecanoyl-*sn*-glycero-3-phosphocholine (PC(17:0/17:0)), 1-heptadecanoyl-2-hydroxy-*sn*-glycero-3-phosphocholine (LPC(17:0)) and 1-palmitoyl-d31-2-oleoyl-*sn*-glycero-3-phosphocholine (PC(16:0/d31/18:1)), purchased from Avanti Polar Lipids, Inc. (Alabaster, AL, USA), and 1,2-Dimyristoyl-*sn*-glycero-3-phospho(choline-d_13_) (PC(14:0/d13)) (Sigma-Aldrich, Steinheim, Germany), tripalmitin-1,1,1-13C3 (TG(16:0/16:0/16:0)-13C3) and trioctanoin-1,1,1-13C3 (TG(8:0/8:0/8:0)-13C3) (Larodan AB, Solna, Sweden). The samples were vortex mixed and incubated on ice for 30 min after which they were centrifuged (9400 × *g*, 3 min, 4 °C). 60 µL from the lower layer of each sample was then transferred to a glass vial with an insert and 60 µL of CHCl_3_:MeOH (2:1, v/v) was added to each sample. The samples were re-randomized and stored at -80 °C until analysis.

Calibration curves using 1-hexadecyl-2-(9Z-octadecenoyl)-sn-glycero-3-phosphocholine (PC(16:0e/18:1(9Z))), 1-(1Z-octadecenyl)-2-(9Z-octadecenoyl)-sn-glycero-3-phosphocholine (PC(18:0p/18:1(9Z))), 1-octadecanoyl-sn-glycero-3-phosphocholine (LPC(18:0)), 1-(1Z-octadecenyl)-2-docosahexaenoyl-*sn*-glycero-3-phosphocholine (PC(18:0p/22:6)) and 1-stearoyl-2-linoleoyl-*sn*-glycerol (DG(18:0/20:4)) from Avanti Polar Lipids, Inc., 1-Palmitoyl-2-Hydroxy-sn-Glycero-3-Phosphatidylcholine (LPC(16:0)) from Larodan, and 1,2,3-Triheptadecanoylglycerol (TG(17:0/17:0/17:0)) and 3β-Hydroxy-5-cholestene 3-linoleate (ChoE(18:2)) from Sigma-Aldrich were prepared prepared to the following concentration levels: 100, 500, 1000, 1500, 2000 and 2500 ng mL^−1^ (in CHCl_3_:MeOH, 2:1, v/v) including 1000 ng mL^-1^ of each internal standard.

The samples were analysed using an ultra-high-performance liquid chromatography quadrupole time-of-flight mass spectrometry method (UHPLC-Q-TOF-MS), which has been presented in detail previously^59^. Briefly, the UHPLC system used in this work was a 1290 Infinity system from Agilent Technologies (Santa Clara, CA, USA). The system was equipped with a multi sampler (maintained at 10 °C), a quaternary solvent manager and a column thermostat (maintained at 50 °C). Separations were performed on an ACQUITY UPLC^®^ BEH C18 column (2.1 mm × 100 mm, particle size 1.7 µm) by Waters (Milford, USA).

The mass spectrometer coupled to the UHPLC was a 6550 iFunnel quadrupole time of flight (Q-TOF) from Agilent Technologies interfaced with a dual jet stream electrospray (dual ESI) ion source. All analyses were performed in positive ion mode and MassHunter B.06.01 (Agilent Technologies) was used for all data acquisition.Quality control was performed throughout the dataset by including blanks, pure standard samples, extracted standard samples and control plasma samples. Relative standard deviations (%RSDs) for retention times and peak areas for lipid standards representing each lipid class in the control plasma samples (*n*= 37) and in the pure standard samples (*n* = 37) were calculated. The %RSDs for the retention times were on average below 0.2% for both the control plasma samples and for the pure standard samples. The peak areas were normalized to the peak area of PC (16:0/d30/18:1) in each showing %RSDs within accepted analytical limits at averages of 13% and 14% for the control plasma samples and for the pure standard samples, respectively. This shows that the method is reliable and repeatable throughout the sample set.

### Data preprocessing

MS data processing was performed using open source software MZmine 2.18^60^. The following steps were applied in the processing: 1.) Crop filtering with a m/z range of 350 – 1700 m/z and a RT range of 2.5 to 21.0 min, 2.) Mass detection with a noise level of 750, 3.) Chromatogram builder with a min time span of 0.08 min, min height of 2250 and a m/z tolerance of 0.006 m/z or 10.0 ppm, 4.) Chromatogram deconvolution using the local minimum search algorithm with a 70% chromatographic threshold, 0.05 min minimum RT range, 5% minimum relative height, 2250 minimum absolute height, a minimum ration of peak top/edge of 1 and a peak duration range of 0.08 -5.0, 5.), Isotopic peak grouper with a m/z tolerance of 5.0 ppm, RT tolerance of 0.05 min, maximum charge of 2 and with the most intense isotope set as the representative isotope, 6.) Peak filter with minimum 12 data points, a FWHM between 0.0 and 0.2, tailing factor between 0.45 and 2.22 and asymmetry factor between 0.40 and 2.50, 7.) Peak list row filter keeping only peak with a minimum of 1 peak in a row, 8.) Join aligner with a m/z tolerance of 0.006 or 10.0 ppm and a weight for of 2, a RT tolerance of 0.1 min and a weight of 1 and with no requirement of charge state or ID and no comparison of isotope pattern, 9.) Peak list row filter with a minimum of 53 peak in a row (= 10% of the samples), 10.) Duplicate peak filter with a m/z tolerance of 0.006 m/z or 10.0 ppm and a RT tolerance of 0.1 min, 11.) Gap filling using the same RT and m/z range gap filler algorithm with an m/z tolerance of 0.006 m/z or 10.0 ppm, 12.) Peak filter using the same parameters as under step 6, 13.) Peak list row filter with a minimum of 265 peak in a row (= 50% of the samples), 14.) Identification of lipids using a custom database search with an m/z tolerance of 0.006 m/z or 10.0 ppm and a RT tolerance of 0.1 min, 15.) Normalization using internal standards (PE (17:0/17:0), SM (d18:1/17:0), Cer (d18:1/17:0), LPC (17:0), TG (16:0/16:0/16:0)-13C3 and PC (16:0/d30/18:1)) using in-house developed R-script, 16.) Data imputation of missing values were done with half of the rows minimum.

### Statistical analysis

All statistical analyses were based on log2-transformed intensity data. The transformed data were mean centered and auto scaled prior to multivariate analysis to improve the global interpretability. The multivariate analysis was done using the PLS Toolbox 8.2.1 (Eigenvector Research Inc., Manson, WA, USA) in MATLAB 2017b (Mathworks, Inc., Natick, MA, USA). PCA of the pre-processed data was initially performed to highlight patterns and to emphasize variation in the dataset. ANOVA-simultaneous component analysis (ASCA) a multivariate extension of ANOVA analysis was performed to allow interpretation of the variation induced by the different factors including age, sex, case, and their interaction. Subsequently, supervised partial least square discriminant analysis (PLS-DA) was performed to discriminate between the samples groups (PT1D/P1Ab/CTR) using combination of various categorical variables (e.g. PT1D vs. CTR, PT1D vs. P1Ab) in a specific age cohort. The area under the receiver operating characteristic (AUROC) were calculated to evaluate the ability of discrimination model to correctly classify the samples. All PLS-DA models were cross-validated by splitting the dataset into test sample subsets for each sub-validation experiment using random subset with ten data splits and twenty repeats. Furthermore, supervised multilevel partial least square discriminant analysis (ML-PLSDA), a multivariate extension of paired t-test was performed to discriminate between matched paired of samples, for instance the before and after seroconversion samples. ML-PLSDA models were validated using double cross model validation (CMV) ^61^ with twenty repeats. The mat lab code for ML-PLSDA is open access and available at http://www.bdagroup.nl/.

For univariate analysis, lipid concentrations were compared using both parametric and non-parametric test as per requirement. Wilcoxon rank-sum test was performed for comparing the two study groups of samples (e.g. PT1D vs. P1Ab) in a specific age cohort assuming data violates heteroscedasticity. For comparison one sample per subject, closest to the age within the time window, has been used in each test. Paired t-test was performed for the matched groups of samples (e.g. before vs. after seroconversion). The resulting nominal p-values were corrected for multiple comparisons using Storey approach^62^ if the number of tests in a study were large (>10) or Benjamin and Hochberg approach^63^ if the number of tests were small (<10). The adjusted p-values < 0.1 (q-values) were considered significantly different among the group of hypotheses tested in a specific age cohort. All of the univariate statistical analyses were computed in MATLAB 2017b using statistical toolbox. The fold difference was calculated by dividing the mean concentration of a lipid species in one group by another, for instance mean concentration in the PT1D by the mean concentration in P1Ab, and then illustrated by heat maps. The individual lipids levels were visualized as box plot using GraphPad Prism 7 (GraphPad Software, La Jolla, CA, USA).

### Data availability

The lipidomics data described in this study are deposited at the MetaboLights database^64^ with the acquisition number (MTBLS620). The associated meta-data were captured using the ISA-creator package available from the MetaboLights. All the data supporting the findings of this study are available from MetaboLights database or from the corresponding authors on reasonable request.

### Ethical approval and informed consent

The ethics and research committee of the participating university and hospital at University of Tampere, Tampere Finland, approved the study protocol. The study was conducted according to the guidelines in the Declaration of Helsinki. All families provided written informed consent for participation in the study.

## Acknowledgments

This work was supported by the JDRF grants 4-1998-274, 4-1999-731 4-2001-435 and special research funds for Oulu, Tampere and Turku University Hospitals in Finland. This work was supported by the Juvenile Diabetes Research Foundation (2-SRA-2014-159-Q-R to M.O.) and the Academy of Finland (Centre of Excellence in Molecular Systems Immunology and Physiology Research – SyMMyS, Decision No. 250114, to M.O. and M.K.).We thank Olli Simell for his contribution in the DIPP study. Anette Untermann for excellent technical support in metabolomics analysis. We also thank Alex Dickens and Partho Sen for useful discussions and insight in relation to this study.

## Author contributions

M.O. and M.K. designed and supervised the study. L.A. and T.S.D. performed metabolomics analysis, which was supervised by T.H. S.L. analyzed the data. H.H., J.I., J.T. and R.V. contributed to the design of the clinical study. S.L. and M.O. wrote the manuscript. All authors critically reviewed and approved the final manuscript.

